# *In vivo* phage display: identification of organ-specific peptides using deep sequencing and differential profiling across tissues

**DOI:** 10.1101/2020.07.01.181974

**Authors:** Karlis Pleiko, Kristina Põšnograjeva, Maarja Haugas, Päärn Paiste, Allan Tobi, Kaarel Kurm, Una Riekstina, Tambet Teesalu

**Affiliations:** Laboratory of Cancer Biology, Institute of Biomedicine and Translational Medicine, University of Tartu, Tartu, Estonia; Faculty of Medicine, University of Latvia, Riga, Latvia; University of Tartu, Department of Geology, Ravila 14A, 50411 Tartu, Estonia; Cancer Research Center, Sanford Burnham Prebys Medical Discovery Institute, La Jolla, California, USA; Center for Nanomedicine and Department of Cell, Molecular and Developmental Biology, University of California, Santa Barbara, Santa Barbara, California, USA

## Abstract

*In vivo* phage display is widely used for identification of organ- or disease-specific homing peptides. However, the current *in vivo* phage biopanning approaches fail to assess biodistribution of specific peptide phages across tissues during the screen, thus necessitating laborious and time-consuming post-screening validation studies on individual peptide phages. Here, we adopted bioinformatics tools used for RNA sequencing for analysis of high throughput sequencing (HTS) data to estimate the representation of individual peptides during biopanning *in vivo*. The data from *in vivo* phage screen were analyzed using differential binding – relative representation of each peptide in the target organ vs. in a panel of control organs. Application of this approach in a model study using low-diversity peptide T7 phage library with spiked-in brain homing phage, demonstrated brain-specific differential binding of brain homing phage and resulted in identification of novel lung and brain specific homing peptides. Our study provides a broadly applicable approach to streamline *in vivo* peptide phage biopanning and to increase its reproducibility and success rate.

**Graphic abstract:** 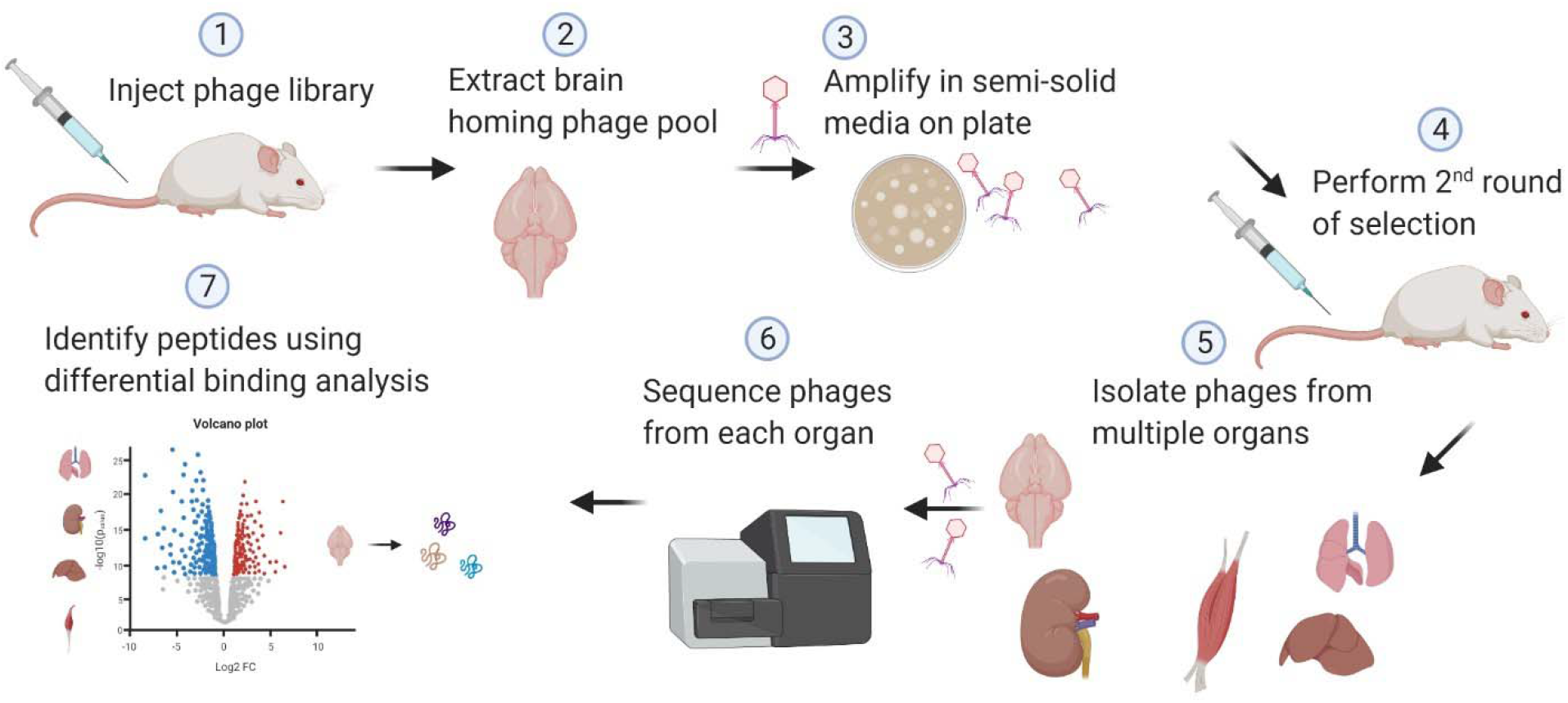

*In vivo* phage display using differential binding approach

## INTRODUCTION

The ability to present foreign peptides as fusions to the nucleocapsid proteins on the surface of bacteriophage was first described by Smith in 1985 (1). Since then, application of the engineered phages for *de novo* peptide ligand discovery has become invaluable and widely applied tool (2). Pasqualini and Ruoslahti introduced *in vivo* phage display in 1996 (3) and adopted the method for identification of peptides that target tumor vasculature in 1998 (4). During *in vivo* biopanning, after short circulation (typically 10 min to 1 h), free phages and phages expressing weakly binding peptide sequences are removed by perfusion. Target tissues, e.g. xenograft tumors, or specific organs, are then collected and tissue-bound phages are rescued by amplification for use in subsequent round of selection (5).

A number of tumor-homing peptides have been identified using *in vivo* phage display. The power of *in vivo* display is particularly well illustrated by identification of a novel class of tumor homing peptides, tumor penetrating peptides, that use a complex multistep mechanism, very hard to envision to be designed rationally, for tumor-specific homing and vascular exit. The prototypic tumor penetrating peptide, iRGD (amino acid sequence: CRGDKGPDC) contains a RGD motif that binds the integrins and a cryptic R/KXXR/K C-end Rule (CendR) motif that is activated by proteolytic cleavage (6). C-terminally exposed CendR motif binds to neuropilin-1 (NRP-1) to trigger endocytic transport and tumor penetration pathway named CendR pathway (7). The use of CendR pathway can improve transport conjugated and coadministered therapeutic compounds into tumors *in vivo* (8). iRGD peptide is clinically developed by CEND Therapeutics Inc. (La Jolla, CA, USA) in phase I clinical trial on pancreatic cancer patients as an add-on treatment in combination with Abraxane and gemcitabine (clinical trial identifier: NCT03517176). Other vascular-homing peptides Cilengitide (sequence: RGDfV) (9), SFITGv6 (sequence: GRCTFRGDLMQLCYPD) (10), CNGRC (sequence: CNGRC) (11), tumor extracellular matrix (ECM) homing peptides DAG (sequence: CDAGRKQKC) (12), ZD2 (sequence: CTVRTSADC) (13), PL1 (sequence: PPRRGLIKLKTS) (14) and PL3 (sequence: AGRGRLVR) (15), IP3 (sequence :CKRDLSRRC) (16), CAR (sequence: CARSKNKDC) (17), and M2 tumor associated macrophage targeting UNO peptide (18) have been selected for possible therapeutic use and reviewed recently (19). Moreover, healthy organs can be specifically targeted with homing peptides. Examples are brain homing peptide CAGALCY (20, 21) and prototypic C-end Rule (CendR) peptide RPARPAR that upon intravenous administration accumulates in lungs and heart (7).

High-throughput sequencing (HTS) of peptide-encoding region of the phage genome can be used to gain insight into the evolution of the peptide landscape during biopanning. HTS has proven particularly useful for binding peptide identification during “low noise” selections on purified individual target proteins (22). However, HTS-based identification of *in vivo* homing peptides remains challenging. One reason is that the peptides that accumulate in intended target tissue can also home to other sites, including healthy organs. Such off-target binding will only be noted when biodistribution of individual phages is assessed. Another complicating factor, especially when using *in vivo* phage display with lysogenic filamentous phages that rely on bacterial cellular machinery for export, is the enrichment of parasitic peptide sequences that increase phage fitness and selection against phages displaying peptides that reduce phage amplification. During M13 phage amplification representation of different peptide phages in a pool can be skewed 10-fold to 100-fold (23). Two approaches can be used to reduce the amplification bias – either avoiding the amplification altogether or performing the amplification in a way that each phage has the same amplification ratio (22, 24). Bioinformatics tools have been developed to prospectively identify parasitic peptide sequences, such as SAROTUP (25) and PhD7Faster (26).

In recent years, a number of studies have been published on HTS-based phage display. Hurwitz et al. developed motif search algorithm that breaks 12-mer peptide sequences into smaller fragments and aligns them with other sequenced fragments using MAFFT software (27). Another motif search algorithm for mapping consensus sequences within the phage display-derived HTS datasets was developed by Rebollo et al. (28). Biopanning Data Bank (BDB) is a large scale effort to combine together data sets acquired from phage display experiments with more than 95 next generation phage display data sets and biopanning data from 1540 published articles (29). PHage Analysis for Selective Targeted PEPtides (PHASTpep) has introduced analysis of both positive and negative selection and sequencing both resulting phage pools. HTS data can be further sorted based on their abundance in each pool (30).

Here we present a statistical analysis-based approach for HTS data mining to determine differential binding of peptide phages across multiple organs during *in vivo* biopanning. Our approach eliminates parasitic peptide sequences in the selected peptide phage pool and minimizes false leads thereby reducing the number of individual peptide phages that need to be used for *in vivo* biodistribution studies.

## MATERIALS AND METHODS

### *In vivo* phage display

For *in vivo* biopanning, we used a model low-diversity (10^5^) CX_7_C T7 phage peptide library with spiked brain-homing peptide (CAGALCY (20, 21)) as a positive control. The library was purified by PEG8000 precipitation and filtered through 0.2 µm syringe filter. For first biopanning round, 100 µL of 5×10^9^ plaque forming units (pfu) CX_7_C T7 library with spiked-in 5×10^4^ pfu CAGALCY in Phosphate Buffered Saline (PBS) was injected intravenously (i.v.) in Balb/C mice. Copy number of CAGALCY phages, 5×10^4^ pfu, was similar to that of individual peptide-phages in the CX_7_C library, as 10^5^ different phages from naïve CX_7_C library were amplified and 5×10^9^ phages were used per injection. The input diversity used is in the same range of a typical peptide phage library after ex vivo preselection step often included prior to in vivo biopanning. After 30 min, the animals were anesthetized and perfused with 10-15 ml of PBS containing 1% BSA (PBS-BSA). The target and control organs (brain, lung, liver, kidney, skeletal muscle) were collected, transferred in 1 mL of LB-1% NP-40, and homogenized on ice using handheld homogenizer with replaceable plastic tips. Phages in tissue lysates were rescued and amplified in *E. coli* strain BLT5403 in semi-solid media, purified by PEG8000 precipitation and filtered through 0.2 μm filter. For 2^nd^ round of selection, 3 brain samples or 3 lung samples from 1^st^ round were mixed and 5×10^9^ pfu in 100 µL were used for injection (n=3 both for lung selection and brain selection) using the same procedure as in 1^st^ cycle. Tissues were dissected and processed identically to 1^st^ cycle. Final libraries were amplified, purified by PEG8000 precipitation and used for sequencing.

The animal procedures were approved by the Committee of Animal Experimentation, Estonian Ministry of Agriculture in accordance with Estonian legislation and Directive 2010/63/EU of the European Parliament and of the Council of 22 September 2010 on the protection of animals used for scientific purposes (permits #48 and #159).

### Determination of phage displayed peptide sequences

Peptide-encoding region of bacteriophage genome was amplified by PCR using Phusion Green Hot Start II High-Fidelity DNA Polymerase (Thermo Scientific, F537L) (reaction volume: 25 µL, list of primers added as Supplementary Table S1). Cycling conditions: denaturation at 98°C for 30 sec., followed by 25 amplification cycles (10 sec at 98°C, 21 sec at 72°C), and final elongation (72°C for 5 min). PCR products were purified using AMPure XP Bead Based Next-Generation Sequencing Cleanup system (Beckmann Coulter, A63881) using 1,8 μL of beads per 1 μL of PCR products. Purified PCR products were quantified using Agilent Bioanalyzer 2100 Instrument using the High sensitivity DNA Kit (Agilent, 5067-4626). Ion Torrent Emulsion PCR and enrichment steps were performed using Ion PGM HiQ View OT2 kit (Life technologies, A29900). HTS was performed using Ion Torrent™ Personal Genome Machine™ (ION-PGM) using Ion PGM HiQ View sequencing kit (Life technologies, A30044) and Ion 316v2 chips (Life Technologies, 448149). The FASTQ sequence files were converted to text files and translated using in-house developed python scripts.

### Peptide data mining

edgeR (31) was used for normalization of peptide count (Input data available as Supplementary Tables S2-S4) in the samples and for statistical analyses. Peptides with <5 reads present in <2 replicate samples were not used for further analysis. Three biological replicates of peptides present in brain or lung samples versus input library were compared for peptide representation analysis. Three biological replicates of peptides present in brain or lung samples versus control organs (liver, kidney, muscle, brain or lung) were compared for differential binding analysis. False discovery rate (FDR) cutoff of 0.05 and logFC cutoff of 2 were used as thresholds for statistical significance. UpSetR (32) package was used to create visualizations of the number of peptides in multiple-organ comparisons. Packages from tidyverse were used to create other sequencing analysis-related graphics (33).

### Peptide phage cloning

Nucleotide sequences coding for BH1 (CLNSNKTNC), BH2 (CWRENKAKC) and LH1 (CGPNRRGDC) peptides were cloned into phage genomic DNA to be expressed on the phage surface as C-terminal fusions to capsid protein. The cloning was performed using complementary oligonucleotides (AATTCTTGCGGCCCGAACCGCCGCGGCGATTGCTA and AGCTTAGCAATCGCCGCGGCGGTTCGGGCCGCAAG for LH1, AATTCTTGCCTGAACA GCAACAAAACCAACTGCTA and AGCTTAGCAGTTGGTTTTGTTGCTGTTCAGGCAAG for BH1, AATTCTTGCTGGCGCGAAAACAAAGCGAAATGCTA and AGCTTAGCATTTC GCTTTGTTTTCGCGCCAGCAAG for BH2). Complementary oligonucleotides were diluted in milli-Q (mQ) water at 100 nM, heated at 95°C for 5 min and slowly cooled to room temperature for annealing. The cloning was performed according to the manufacturer’s protocol using the T7Select® 415-1 Cloning Kit (Millipore, 70015-3) with 1 μL of annealed oligonucleotides.

### Intra-animal validation of candidate phages

HTS play-off technique allows comparative internally-controlled auditioning of tissue selectivity of systemic candidate peptide phages. Phages expressing a BH1, BH2, LH1, CAGALCY or G7 (CGGGGGGGC) peptides were amplified in *E. coli* strain BLT5403, purified by PEG8000 precipitation, followed by cesium chloride gradient centrifugation, and dialyzed against PBS. The phages were mixed in equimolar ratio and 1×10^10^ pfu of phage mix was injected i.v. into Balb/C mice (n=3). After 30 min, the animals were anesthetized and perfused with PBS-BSA. The target and control organs (brain, lung, liver, kidney, skeletal muscle) were collected, the tissues were homogenized and phages in tissue lysates were amplified and purified by PEG8000 precipitation. The distribution of phages between target and control organs was determined by HTS. Fold change was calculated as follows:

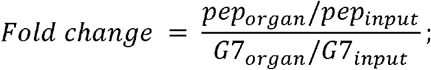

Where:

pep_organ_ - % of reads coming from the peptide of interest in particular organ sequencing dataset;

pep_input_ - % of reads coming from the peptide of interest in input library sequencing dataset;

G7_organ_ - % of reads coming from G7 peptide in particular organ sequencing dataset

G7_input_ - % of reads coming from G7 peptide in input library sequencing dataset

### Inter-animal validation of candidate phages

Phage biodistribution studies by titering are widely used to study the biodistribution of the single peptide phages in individual animals. For titering-based biodistribution studies, 5×10^9^ pfu of phage expressing BH1, BH2, LH1, CAGALCY or G7 peptide was injected i.v. into Balb/C mice (n=3). After 60 min, the animals were anesthetized and perfused with PBS-BSA. The target and control organs (brain, lung, liver, kidney, skeletal muscle) were collected, homogenized, and the amount of phage in tissue lysates was determined by titering using *E*.*coli* strain BLT5615. Fold change was calculated as follows:

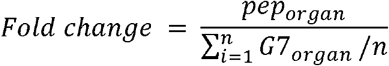

Where:

n – number of animals per group (n=3 in this case);

pep_organ_ – pfu/mg titer from peptide coding phage of interest in particular organ;

G7_organ_ – pfu/mg titer from G7 coding phage in the particular organ

### Database search for target-unrelated peptides

All sequences that were identified using differential binding approach were searched in SAROTUP (http://i.uestc.edu.cn/sarotup3/) (25) to identify target-unrelated peptides (TUPScan: http://i.uestc.edu.cn/sarotup3/cgi-bin/TUPScan.pl), peptides containing sequences known to cause phages to grow faster (PhDFaster 2.0: http://i.uestc.edu.cn/sarotup3/cgi-bin/PhD7Faster.pl) and already known sequences in BDB (MimoScan: http://i.uestc.edu.cn/sarotup3/cgibin/MimoScan.pl).

### Synthesis and functionalization of silver nanoparticles

For peptide biodistribution visualization purpose, Ag nanoparticle labeled peptides were used. The synthesis and surface functionalization of silver nanoparticles (AgNPs) was based on previously published methods (21, 34, 35) with minor modifications. Biotin-Ahx-LGDPNS-CAGALCY-OH (CAGALCY), biotin-Ahx-CLNSNKTNC-OH (BH1), biotin-Ahx-CWRENKAKC-OH (BH2) (TAG Copenhagen A/S, Frederiksberg, Denmark) were used as targeting moieties. Briefly, ultrapure mQ water (0.5 L; resistivity 18 MΩ cm^−1^) in a foil-wrapped flask cleaned with a piranha solution (H_2_SO_4_/H_2_O_2_) was heated to 65°C with stirring. For isotopic barcoding, ^107^AgNO_3_ or ^109^AgNO_3_ (50.4 mg; Isoflex USA, San Fransisco, CA, USA) pre-dissolved in 1 ml of mQ water were added to the flask. Next, trisodium citrate hydrate (200 mg; #25114, Sigma-Aldrich Co., LLC) and tannic acid (1.2 mg; #403040, Sigma-Aldrich Co., LLC) were pre-dissolved in mQ water (20 mL) and added to the vessel. The solution was heated at 65°C for 3 min, followed by stirring on a hot plate at 250°C in the dark for 27 min. NeutrAvidin (NA; #31055.

Thermo Scientific Inc., Washington, USA) was modified with an OPSS-PEG(5K)-SCM linker (OPSS; JenKem Technology USA Inc., Texas, USA) according to the procedure described by Braun *et al*. (34). Subsequently, NeutrAvidin-OPSS (3.9 mL, 2.9 mg/mL) was added to the core AgNPs (500 mL). After 2 min, 4-morpholineethanesulfonic acid hemisodium salt (5 mL, 0.5 M in mQ water; #M0164, Sigma-Aldrich Co., LLC) was added. The pH of the solution was adjusted to 6.0 with 1 N NaOH, and the solution was incubated in a water bath at 37°C for 24 h. The solution was brought to room temperature (RT) and 10X phosphate buffered saline (50 mL; PBS; Naxo OÜ, Tartumaa, Estonia) was added, followed by Tween-20 (250 µL; #P9416, Sigma-Aldrich Co., LLC). The solution was centrifuged at 17,500 × g for 20 min (4°C), the supernatant was removed, and the particles were resuspended in PBST (0.005% Tween-20 in PBS). Next, tris(2-carboxyethyl)phosphine hydrochloride solution (TCEP; #646547, Sigma-Aldrich Co., LLC) was added to a final concentration of 1 mM, followed by incubation at RT for 30-min. Then, lipoic acid-PEG(1k)-NH_2_ (#PG2-AMLA-1k, Nanocs Inc., New York, USA) was added to a final concentration of 5 µM, and the mixture was incubated at RT for 2.5 h. The solution was centrifuged at 17,500 × g at 4°C for 20 min, the supernatant was removed and the particles were resuspended in PBST to half of the initial volume. The AgNP solution was filtered through a 0.45 µm filter and stored at 4°C in dark.

CF555 dye (#92130, Biotium Inc., California, USA) with an NHS group was coupled to the NH_2_ groups on the AgNPs. CF555-NHS (5 µL, 2 mM) in dimethyl sulfoxide (DMSO; Sigma-Aldrich Co., LLC) was added to AgNPs (500 µL), followed by overnight incubation at 4°C. The particles were washed 3 times by centrifugation at 17,500 × g at 4°C for 20 min, followed by resuspension of the particles in PBST. Next, biotinylated peptides were coupled to the particles by adding peptide (10 µL, 2 mM in mQ water) to AgNPs (500 µL), followed by incubation at RT for 30 min. The AgNPs were washed, 0.2 µm filtered and stored at 4°C in the dark.

### Mapping of AgNP biodistribution using Laser Ablation Inductively Coupled Plasma Mass Spectrometry (LA-ICP-MS)

The ^107^AgNPs (functionalized with a peptide (CAGALCY, BH1 or BH2) and control ^109^AgNPs (blocked with biotin) were mixed in equimolar ratio and i.v. injected into mice. In total, 200 μl of AgNPs (OD_415_=100) was injected per mouse. After 3 h of circulation, the mice were anesthetized and perfused with PBS. The brain and liver were collected, frozen in OCT, sectioned at 30 μm on Superfrost^+^ slides, and dried in a vacuum desiccator.

LA-ICP-MS analysis of the tissue samples was performed using Agilent 8800 ICP-MS/MS coupled to a Cetac LSX-213 G2+ laser ablation unit equipped with HelEx II ablation cell and connected using ARIS (Aerosol Rapid Introduction System) sample introduction system. The system was tuned using NIST 610 glass. Ablation was performed as line scans using 65 µm square spot, scan speed of 260 µm/s, 20 Hz and fluence of 13.5J/cm2. 5 parallel ablations with 65 µm spacing were performed on each tissue. ICP-MS was operated in single quad mode. Data collection was performed in TRA mode with dwell times of 9.5 ms on mass 13C and 14 ms on mass ^107^Ag and ^109^Ag corresponding to a total duty cycle of 50ms. Data reduction was performed using Iolite v3.62. Median with MAD error 2 S.D. outlier reject was used for data selection.

## RESULTS

### Identification of candidate brain and lung homing peptides using *in vivo* peptide phage display with low complexity library

Low-diversity naïve CX_7_C T7 phage peptide library with spiked-in brain-homing peptide CAGALCY was used as the starting library for *in vivo* phage display screens. For round 1 of biopanning, the starting library was injected into mice (n=3) and after 30 min of circulation, the mice were perfused to remove free and weakly bound background phages. Target (lung, brain) and control organs (liver, kidney, skeletal muscle) were collected, phages from these organs were amplified and the samples were subjected to Ion-Torrent HTS. Starting library was sequenced in 3 technical replicates. Phage pools recovered from lung and brain were used for two separate 2^nd^ biopanning rounds (n=3 for each). After round 2, all control and target organs were collected, and phage pools were sequenced (Fig. 1). Sequencing was also performed for input libraries.

**Fig. 1.**
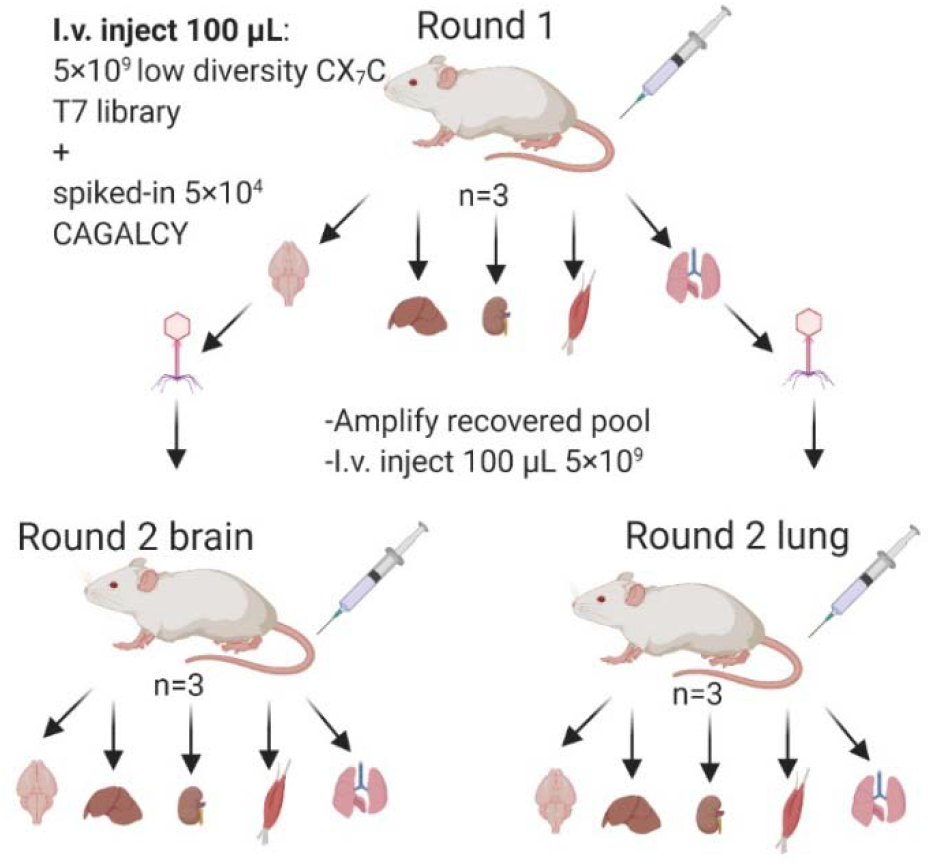
Flow of in vivo phage biopanning. In round 1, 100 µL of 5×10^9^ pfu collapsed CX_7_C T7 phage library and 5×10^4^ pfu of CAGALCY coding phage was injected in tail vein of mice (n=3). Target (lung and brain) and control organs (kidney, liver, muscle) were collected and phages were rescued by amplification in E. coli. For round 2, 5×10^9^pfu of amplified libraries were injected in mice for further enrichment of lung (n=3) or brain (n=3) selective phages. All phage pools were subjected to HTS of the variable peptide encoding region of the phage genome.

The sequences of peptides expressed on the surface of phages recovered from target and control tissues were derived from the DNA sequencing data. The representation of reliable peptides, defined as peptides present in all replicates, was determined for three technical replicates of round 1 input library and for three biological replicates of phages amplified from organs (Fig. 2). The number of reliable peptides for input libraries is the same as the depth of sequencing since these samples were prepared as technical replicates from one input sample. At the sequencing depth used, reliable peptides in round 1 input library made up ∼30% of all peptides. Phage pools recovered from brain displayed smaller fraction of reliable peptides (∼10%). Percentage of reliable peptides in brain was the lowest, while in other organs percentage of reliable peptides from round 1 and round 2 brain samples were comparable to the one in technical replicates. Round 2 lung samples contained a smaller fraction of reliable peptides.

**Fig. 2.**
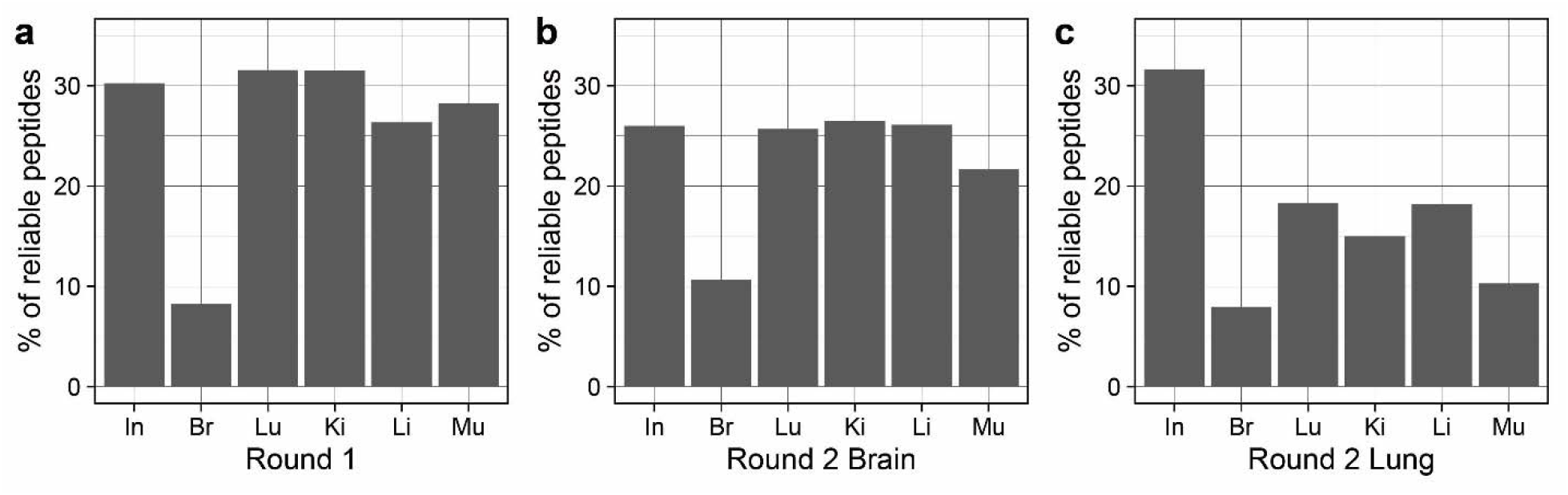
Percentage of reliable peptides during in vivo selection rounds 1 and 2. The graphs represent percentage of reliable peptides (defined as being present in n=3 replicates for each sample) in mice dosed with naïve **(a)**, round 1 brain-selected **(b)**, or round 1 lung-selected libraries **(c)**. Input libraries represent technical variability between technical replicates (n=3) whereas organ-rescued libraries represent biological variability (n=3).

### Differential binding approach identifies organ-selective peptides

Organ selectivity of peptides was determined using differential binding approach, with statistical comparisons performed for each control organ (lung, kidney, liver, and muscle in the case the brain is the target organ) and for the target organ (brain) by using logFC>2 and FDR <0.05 as defined cutoffs. Each peptide that appeared brain-selective based on these criteria, was analyzed further. Peptides from each comparison (e.g., lung-brain, kidney-brain, liver-brain and muscle-brain) were then cross-checked with all identified peptides in other comparisons. Target organ selective peptides were defined as being present in all comparisons. Peptides that were selective only when compared to one or two control organs were discarded (Fig. 3).

**Fig. 3.**
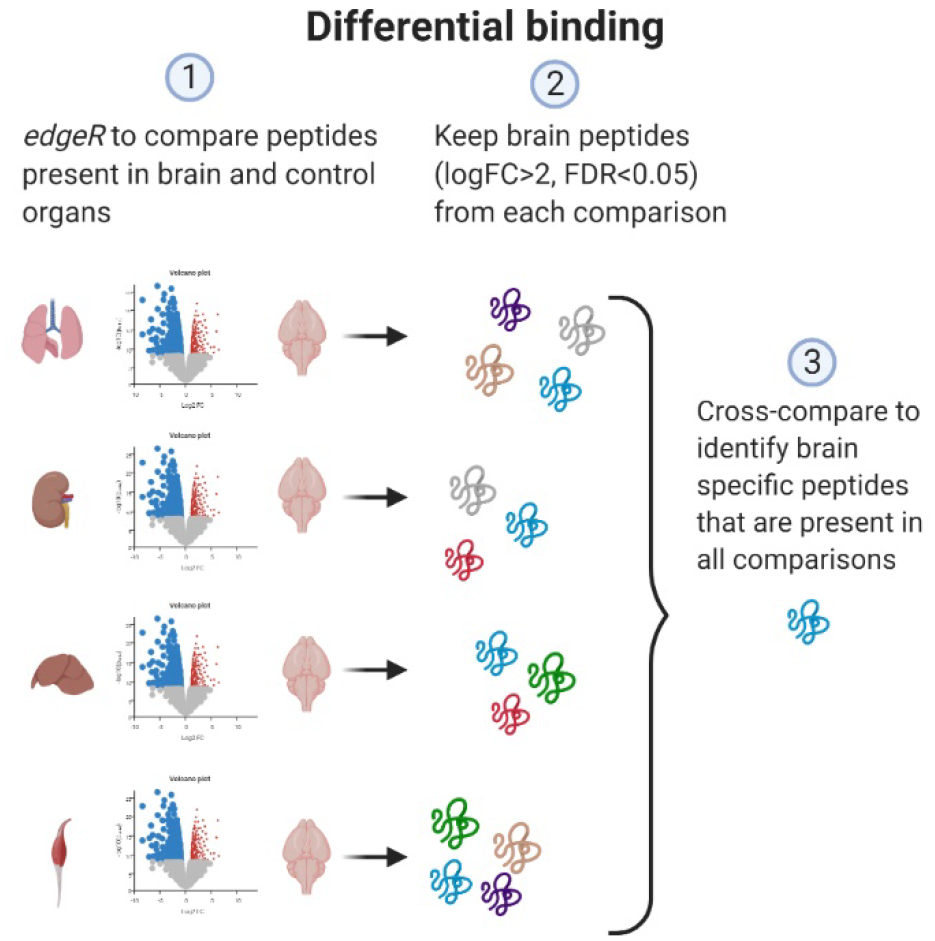
Differential binding analysis. HTS data from target organ (brain or lung) were compared to all control organs individually. Using edgeR, logFC > 2 and FDR <0.05 were applied as cutoff to identify peptides that are statistically significantly over-represented in brain. Peptides from each comparison were cross-compared to identify hits that appear in all comparisons as brain selective.

Landscapes of peptides displayed on phages recovered from brain samples in round 1 of biopanning were compared to those in the kidney, liver, muscle and lung. This analysis yielded a single brain-selective peptide when compared against the muscle sample. In round 2 of selection, 17 peptides were selective for the brain compared to the muscle, 9 compared to the liver, 7 compared to the kidney and 1 peptide (CAGALCY, known brain homing peptide spiked in the starting library) compared to the lung. CAGALCY was brain selective when compared to all control organs and 6 peptides appeared brain-selective when compared against kidney, liver and muscle (Fig. 4a). Based on logFC difference against control organs, BH1 (CLNSNKTNC) and BH2 (CWRENKAKC) were selected for individual testing as the most promising candidate brain homing peptides (Supplementary Table S5).

**Fig. 4.**
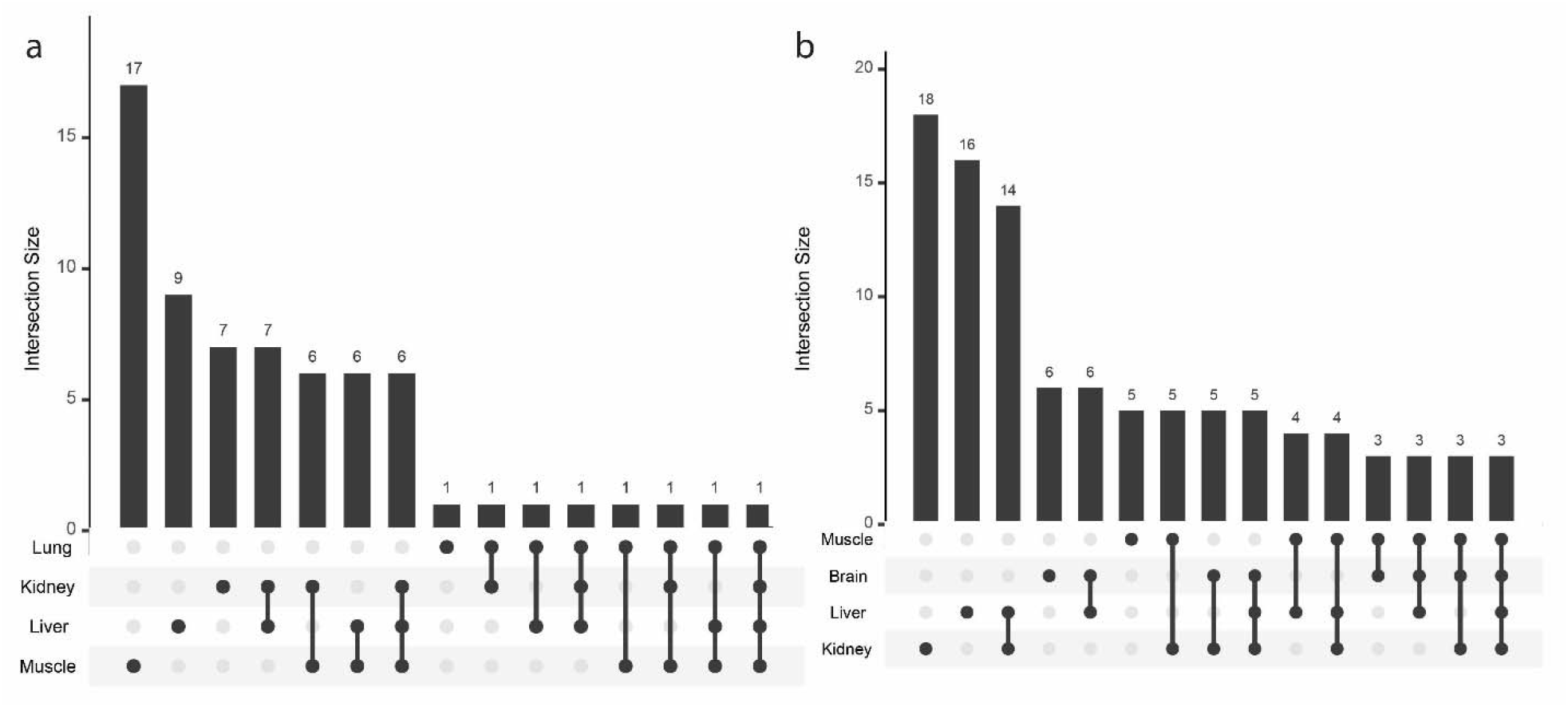
Multiorgan comparison of selected phage pools to identify organ specific homing peptides. Statistically significant (FDR < 0.05) peptides homing to the brain (a) and the lungs (b) compared against control organs (logFC > 2, FDR < 0.05) and number of peptides that overlap between different comparisons. One peptide (CAGALCY) is fully brain selective and 3 peptides lung selective when compared to other organs (Supplementary Tables S5 and S6).

Differential binding analysis for lung-selective peptides in mice dosed with naïve library (round 1) yielded no lung selective peptides when compared to any other control organ. In contrast, round 2 yielded 18 lung selective peptides when the dataset was compared against the kidney, 16 peptides when compared against the liver, 6 when compared against the brain and 5 when compared against the muscle. Three peptides were present in all comparisons as lung-selective peptides (Fig. 4b). Notably, among 20 peptides that were lung selective compared to at least one organ, 8 contained the CX_5_RGDC motif and 7 ended with C-terminal arginine residue (Supplementary Table S6). All three of the identified lung-specific peptides contained the RGDC motif. LH1 (CGPNRRGDC) was selected for further testing, as this peptide had the highest logFC difference toward lung when compared to the brain (logFC = 4.3) and liver (logFC = 3.9).

Enrichment analysis is the most commonly used approach to identify peptides in phage display. We next analyzed enrichment of reliable peptides in target organs in round 1 vs. round 2. Theoretically, the peptides that are most enriched, are most likely to reach the target and possess desired binding properties. We used gating logFC >2 and FDR < 0.05 between round 1 input library and phages recovered from the target organs in rounds 1 and 2. As a threshold, we decided to apply the criteria used for differential binding analysis. After first round of selection, this approach resulted in identification of 4 peptides that are enriched in the lung and no brain-enriched peptides. After round 2, 16 peptides were found enriched in the brain (Supplementary Table S7) and 37 peptides in the lung (Supplementary Table S8).

One of 4 peptides enriched in the lungs after round 1 was an R-ending peptide, KTGARKR; the rest of sequences contained KLAALE phage vector-derived sequence. After round 2, among the 16 peptides enriched in the brain, CAGALCY was the most enriched sequence, followed by CVRLNKVRC (EN-2) and CTVTNKVRC (EN-3). EN-2 sequence in differential binding dataset shows selectivity only against the muscle and EN-3 sequence is selective for the brain when compared against the liver, kidney and muscle. Both BH1 and BH2 sequences that were identified using differential binding approach were present on the list as number 8 and number 4 accordingly. Of 37 peptides identified through enrichment analysis in lung, LH1 was 16th most enriched sequence.

### Selected peptide phages show expected *in vivo* tropism in playoff analysis

In an earlier study, we developed a playoff analysis for the validation of candidate homing peptides (5). The method is based on medium-throughput internally controlled auditioning of mixtures of selected peptide phages. An equimolar mix of CAGALCY, BH1, BH2, LH1 and G7 control phages was i.v. injected into mice (n=3). Control organs (kidney, liver, muscle) and target organs (brain, lung) were collected and phage pools in each organ were sequenced using HTS. The input mixture was also sequenced. After normalization against representation in the input, fold change over G7 phage was calculated for each peptide phage in each organ (Fig. 5).

**Fig. 5.**
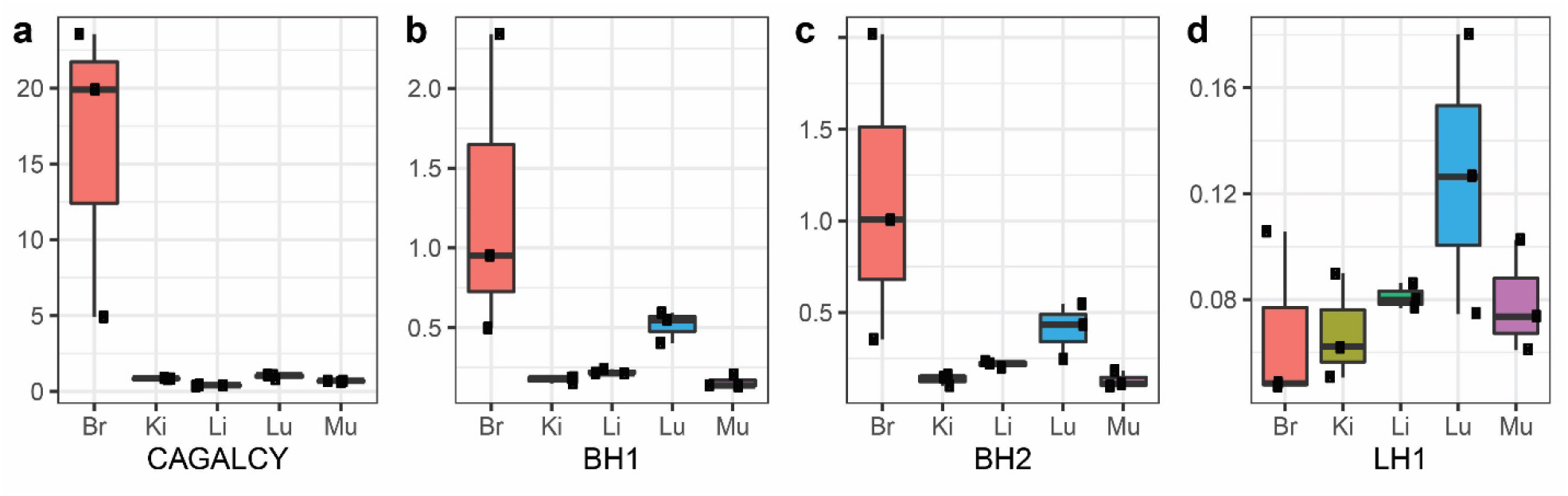
Validation of homing specificity by in vivo peptide phage playoff and HTS. Individual peptide phages (G7, CAGALCY, BH1, BH2, LH1) were mixed at equimolar ratio and injected in BalbC mice (n=3). Results are expressed as fold change over G7 control phage. CAGALCY (a), BH1(b), BH2(c), LH1(d). Lower and upper hinges correspond to first and third quartiles, whiskers represent 1.5*IQR (inter-quartile range), middle line represents the median.

The phage displaying known brain-homing peptide CAGALCY showed the highest fold-change over the G7 control phage (Fig. 5a), both in brain homing and selectivity. Also, BH1 and BH2 showed preferential accumulation in the brain (Fig. 5b-c) and LH1 showed tropism for the lungs (Fig. 5d).

### *In vivo* homing of individual peptide phages

Biodistribution of peptide phages may be affected by other phages in the mix, with the bystander effect best characterized for tumor penetrating CendR peptides (36). Therefore, we decided to study biodistribution of individual candidate organ specific peptide phages.

Individual phage clones displaying CAGALCY, BH1, BH2 and LH1 were i.v. injected in mice (n=3 for each peptide sequence). After 30 min circulation, the organs (brain, lung, liver, kidney, muscle) were collected and phage titer in each organ was determined. For each peptide phage, tissue distribution was expressed as fold change over control G7 phage (Fig. 6).

**Fig. 6.**
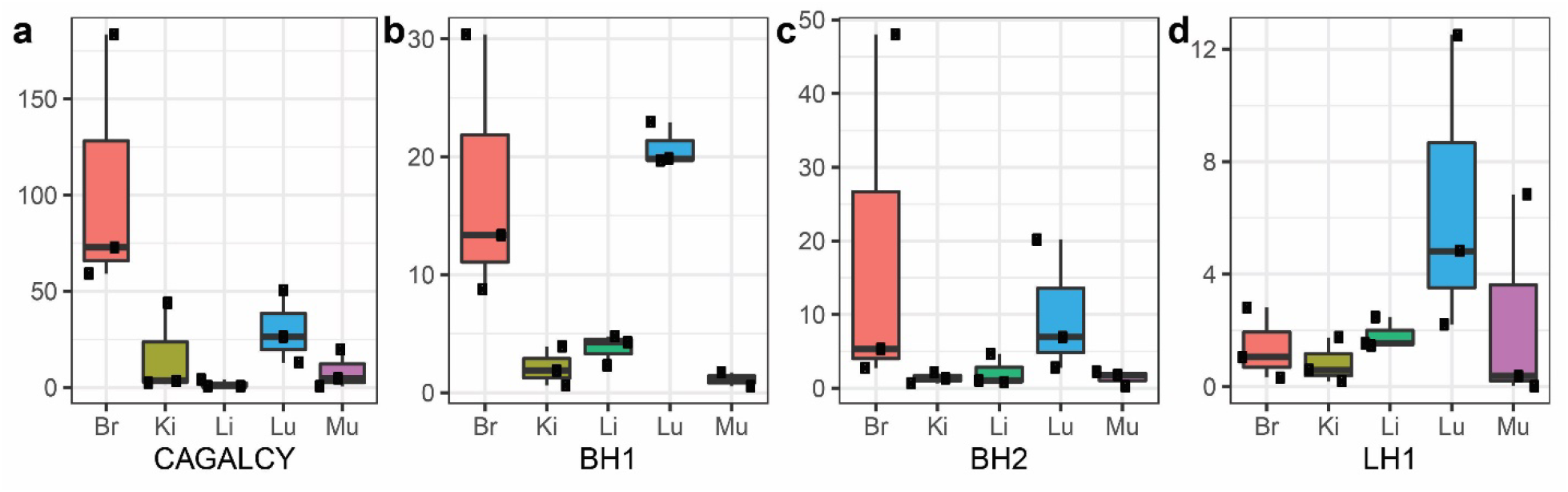
In vivo biodistribution of individual peptide phages. Individual peptide- displaying phages (CAGALCY, BH1, BH2, LH1, G7) were injected in mice (n=3 for each peptide). Organs (brain, lung, liver, kidney, muscle) were collected and phages bound to the organ were titered. Fold change (y axis) was calculated over the titer of G7 phage. CAGALCY (a) homes to brain. BH1(b) and BH2 (c) are selective towards brain when compared to other organs except lung. LH1 (d) shows selectivity towards lung. Lower and upper hinges correspond to first and third quartiles, whiskers represent 1.5*IQR (inter-quartile range), middle line represents the median.

In the brain, CAGALCY-displaying phages showed ∼ 65-fold overrepresentation compared to G7 phage, and the phages displaying newly identified brain homing peptides BH1 and BH2 showed ∼20-fold overrepresentation. Second highest difference for BH1 and BH2 was observed in the lung. This was expected as both sequences did not show brain selectivity when compared to lung using differential binding approach.

Finally, we investigated whether BH1, BH2 or LH1 could be target-unrelated peptides, i.e. peptides that are enriched for reasons other than target recognition. TUPScan did not identify peptides BH1, BH2 or LH1 as target-unrelated peptide sequences. Furthermore, PhDFaster did not detect peptides BH1, BH2, LH1 as sequences that could cause phages to be amplified faster than other clones. MimoScan confirmed that BH1, BH2 and LH1 have not been previously identified in other phage display datasets that are included in BDB.

### Brain-selective peptide functionalization increases delivery of AgNPs to the brain

Effect of brain homing peptides on biodistribution of synthetic nanoparticles was tested using peptide-coated isotopically barcoded AgNPs. Biotinylated CAGALCY, BH1 and BH2 peptides were coated on neutravidin-^107^AgNPs and biotin-blocked neutravidin^109^AgNPs were used as a control. Equimolar cocktail of targeted and control ^107/109^AgNPs was injected i.v. in mice; after 3 h circulation, animals were sacrificed, tissues isolated and AgNPs mapped in brain and liver sections using LA-ICP MS. For all peptide-AgNPs, the liver ratio of ^107/109^AgNPs (∼1) was similar to the ratio of the particles in the input mixture. In the brain, the ratiometric ^107/109^AgNP analysis was performed by quantitation of ^107^Ag and ^109^Ag in five 5 laser ablation paths per cryosection (Fig. 7). The ^107/109^AgNP ratio was 1.62 (95% CI: 1.62 ± 0.0719) for the mice injected with CAGALCY-^107^AgNPs, 1,18 (95% CI: 1.18 ± 0.0331) for BH1-^107^AgNPs, and 1.04 (95% CI: 1.04 ± 0.027) for BH2-^107^AgNPs.

**Fig. 7.**
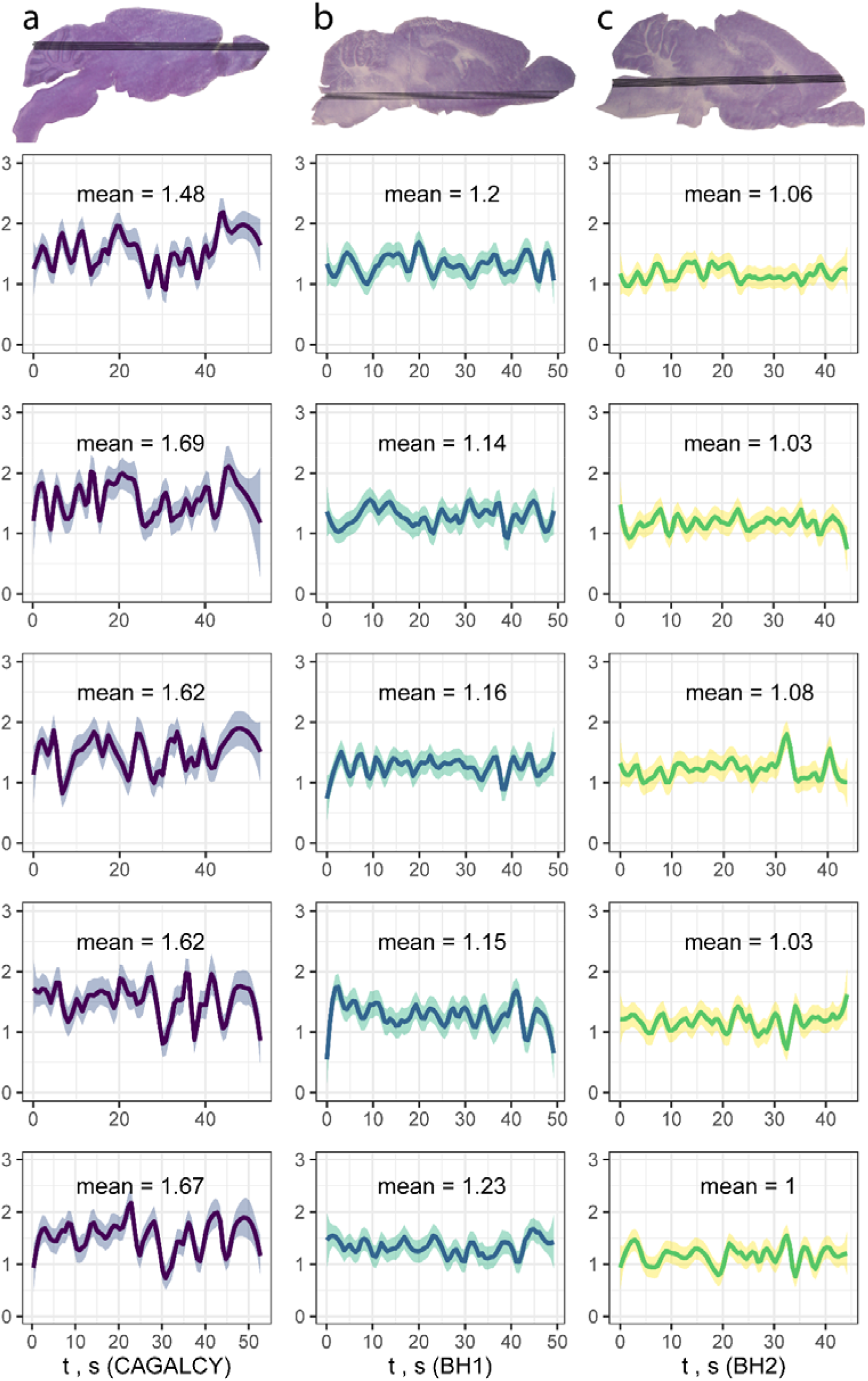
Ratiometric *in vivo* auditioning of brain selectivity of peptide-AgNPs. CAGALCY (1^st^ column), BH1 (2^nd^ column), BH2 (3^rd^ column) were coated on ^107^AgNPs, mixed with equimolar biotin-blocked ^109^AgNPs, and injected i.v. to mice. Brains were isolated after 3 h circulation, ^107^/^109^Ag ratio (plotted on y axis) was determined using LA-ICP-MS over 5 parallel ablation paths (shown as individual graphs; x axis represents the ratio over the length, expressed as acquisition time, of the ablation line). Mean ^107^/^109^Ag ratio was 1.62 (95% CI 1.55 to 1.69) for CAGALCY, 1.18 (95% CI 1.15 to 1.21) for BH1 and 1.04 (95% CI 1.01 to 1.07) for BH2.

## DISCUSSION

*In vivo* peptide phage display is a powerful technology that allows unbiased agnostic mapping of systemically accessible vascular heterogeneity in diseased and normal tissues and development of peptide affinity ligands for precision delivery of drugs, imaging agents and nanoparticles. However, the technology is laborious, and subject to biases and artefacts due to low stringency and reproducibility, and lack of validated tools for identification of the most avid binders in selected phage pools. Availability of powerful and affordable HTS platforms has made it feasible to determine homing peptide phage landscapes in target and control organs. In the current study, we developed an approach for identification of target-selective homing peptides based solely on *in vivo* display HTS data. Application of this approach enables faster and more reliable homing peptide discovery than provided by the current methods. In a study using a peptide T7 phage library with spiked-in brain homing phage demonstrated brain-specific differential binding of brain homing phage and resulted in identification of novel lung and brain selective homing peptides.

Our HTS-based screening addresses the main shortcoming in HTS-based phage biopanning – excessive reliance on enrichment of peptide phages at the target site. Enrichment alone does not provide full indication of selectivity of the peptide, as representation of peptide phages at the target site is also influenced by non-specific promiscuous interactions and peptide-dependent variation in the phage amplification rate (30). Our approach takes into account the statistically weighted enrichment for each peptide in the pool in the context of representation across target and nontarget sites. The method was inspired by published analysis of RNA sequencing data (37) and availability of computational tools such as *edgeR* that allow the use of quasi-likelihood F-tests and correct the results taking into account multiple comparisons (31, 38).

To validate our approach to *in vivo* homing peptide identification, we decided to perform a model screen focused on the enrichment and tissue distribution analyses of a known brain-selective peptide, CAGALCY (20, 21), spiked into a low complexity peptide phage library. We showed that by using the differential binding approach we can not only select for CAGALCY as a brain homing peptide, which would be possible by conventional enrichment analysis, but also profile its biodistribution and selectivity across organs. The diversity of the peptide library used in the model screen, ∼1×10^5^pfu, is ∼4 orders of magnitude lower than the diversity of the naïve CX_7_C library. This is similar to typical diversity of the phage library after preselection *ex vivo*, or on cultured cells, steps often used to enrich for binding peptides prior to *in vivo* biopanning (5). Remarkably, even with the low complexity peptide library used here, the differential binding approach resulted in identification of several lung and brain selective homing peptides. Individual phage homing and play-off studies confirmed that newly identified BH1, BH2, and LH1 peptide phages possess the intended systemic tropism. Complementary *in vivo* ratiometric studies with isotopically barcoded AgNPs confirmed the brain homing of CAGALCY-AgNPs (39) and showed that surface BH1 and BH2 peptide functionalization had a positive effect on AgNP brain delivery.

Similar strategies for novel peptide identification using phage screens that take advantage of statistical methods have already proven effective for HTS data in *in vitro* settings (30). HTS-based peptide differential profiling can be used for systematic identification of homing peptides and possibly other phage-displayed targeting ligands (e.g. single chain antibodies, affibodies). Importantly, the differential binding approach developed here avoids the bias introduced during a phage amplification step between phage display rounds - one of the main drawbacks of the phage display (40).

It should be possible to develop affinity ligands targeting vascular ZIP codes – systemically accessible vascular heterogeneity across normal and diseased tissues (41) following a similar process. Variations on the theme to be explored in follow-up studies include additional tissue fractionation steps for profiling of cell type and extracellular matrix specific peptides in normal organs and in pathological conditions such as tumors.

In summary, differential binding-based HTS *in vivo* phage display developed in the current study is likely to render identification of vascular targeting peptides more reliable and consistent facilitating progress towards development of affinity guided smart drugs and contrast agents.

## DATA AVAILABILITY

R script for differential binding analysis and generation of figures is available at https://github.com/KarlisPleiko/diff_phage.

## ACKNOWLEDGMENTS

We thank Prof. Erkki Ruoslahti for reading and constructive criticism of the manuscript.

## Author contributions

K.P., K.P., M.H. designed and carried out experiments, analyzed the data, composed the figures and wrote the manuscript. U.R. designed the differential binding experiments. K.K. performed sequencing. P.P. performed LA-ICP-MS analysis. A.T. prepared ^107^Ag NPs and ^109^Ag NPs and performed the peptide functionalization of the AgNPs. T.T. conceived and supervised the project, analyzed the data and wrote the manuscript.

Graphic abstract, Fig.1 and Fig. 3 were created using BioRender.com.

